# Quantitative prediction of conditional vulnerabilities in regulatory and metabolic networks of *Mycobacterium tuberculosis*

**DOI:** 10.1101/2021.01.29.428876

**Authors:** Selva Rupa Christinal Immanuel, Mario L. Arrieta-Ortiz, Rene A. Ruiz, Min Pan, Adrian Lopez Garcia de Lomana, Eliza J. R. Peterson, Nitin S. Baliga

## Abstract

The ability of *Mycobacterium tuberculosis* (Mtb) to adopt heterogeneous physiological states, underlies its success in evading the immune system and tolerating antibiotic killing. Drug tolerant phenotypes are a major reason why the tuberculosis (TB) mortality rate is so high, with over 1.8 million deaths annually. To develop new TB therapeutics that better treat the infection (faster and more completely), a systems-level approach is needed to reveal the complexity of network-based adaptations of Mtb. Here, we report a new predictive model called PRIME (**P**henotype of **R**egulatory influences **I**ntegrated with **M**etabolism and **E**nvironment) to uncover environment-specific vulnerabilities within the regulatory and metabolic networks of Mtb. Through extensive performance evaluations using genome-wide fitness screens, we demonstrate that PRIME makes mechanistically accurate predictions of context-specific vulnerabilities within the integrated regulatory and metabolic networks of Mtb, accurately rank-ordering targets for potentiating treatment with frontline drugs.

## INTRODUCTION

*Mycobacterium tuberculosis* (Mtb) kills more people than any other microbe, and it has thus far resisted every attempt to bring the pandemic under control. Part of the pathogen’s success is its ability to diversify itself phenotypically and survive both host and drug bactericidal action^1–3^. Phenotypic heterogeneity (both stochastically and environmentally induced) seems to be an intrinsic characteristic of the pathogen and a major reason why standard chemotherapy of tuberculosis (TB) requires 6 months of treatment, and 5% are not cured even then^4,5^. To develop better interventions that account for pathogen heterogeneity, we need to identify the most important factors (e.g., transcriptional regulators) that create variation as well as the downstream effectors (e.g., regulatory target genes) that mediate drug tolerance.

Metabolic activity undoubtedly contributes to Mtb phenotypic heterogeneity and antibiotic tolerance. For example, changes in metabolism can affect the amount of drug target present^6^, the ability to generate toxic products^7^, and the efflux of antibiotics^8^. Mtb alters its growth and metabolism in response to stressful conditions through regulatory programs primarily encoded at the transcriptional level. Indeed, modeling host-related stresses *in vitro* produces large transcriptional changes in Mtb, particularly in metabolic pathways; consistently ~25% of differentially expressed genes are metabolic genes from hypoxic (GSE116353)^9^, acidic pH (GSE165514), or nutrient limited (GSE165673) conditions. To develop effective antibiotic regimens, we need to understand at a systems- and mechanistic-level how specific regulatory mechanisms conditionally activate and repress genes to redirect flux through metabolic networks to generate and support drug tolerant phenotypes. This mechanistic understanding will uncover new vulnerabilities in Mtb’s regulatory and metabolic networks that can be rationally targeted in new drug regimens to achieve faster and complete clearance of the pathogen.

Previously, approaches to model the influence of transcriptional regulation on metabolism have used boolean logic (Regulatory Flux Balance Analysis – rFBA)^10^, protein-DNA (P-D) interactions (Probabilistic Regulation of Metabolism – PROM)^11,12^, and regression-based regulatory influences (Integrated Deduced REgulation And Metabolism – IDREAM)^13^ to predict how transcriptional regulation of enzyme-coding genes modulates flux through their catalyzed reactions. Briefly, rFBA models the influence of transcriptional regulation on metabolism using boolean “on or off” states of metabolic genes, depending on the expression level of the transcription factor (TF) and its implicated role as a putative activator or repressor of that gene. The extensive manual curation required to develop rFBA and its inability to model TF activity as a continuous (i.e., not boolean) function greatly limits its application and accuracy. In contrast, PROM outperformed^11^ rFBA by using a probabilistic approach to model the regulation of a metabolic gene by a TF using a compendium of transcriptome profiles to calculate probabilities. However, PROM is limited in that it relies on a P-D interaction map for the regulatory network. P-D interactions are typically generated in a limited set of conditions by using an overexpressed TF as a bait to enrich and locate its genome-wide binding locations. P-D interactions are fraught with false positives (due to TF overexpression) and false negatives (due to lack of context for TF regulation across environmental conditions). Notwithstanding these caveats, PROM was useful in uncovering the mechanism by which pretomanid potentiates bedaquiline action on Mtb by disrupting a regulatory network that confers tolerance to the recently approved FDA drug^14^. A third model, IDREAM addressed the shortcoming of using P-D interactions in PROM by constraining flux using TF regulatory influences from a predictive systems-scale **e**nvironment and **g**ene **r**egulatory **i**nfluence **n**etwork (EGRIN) model. An EGRIN model is inferred in two steps using (a) cMonkey, which identifies the specific context in which subsets of genes are co-regulated (biclusters) by a conserved regulatory mechanism(s); and (b) Inferelator, which predicts TFs and environmental factors that causally influence the differential expression of genes within those biclusters^15–17^. By integrating confidence scores for EGRIN-inferred regulatory influences, IDREAM achieved significantly better performance than rFBA and PROM in predicting synthetic lethal interactions between TFs and metabolic genes in yeast^13^. However, IDREAM does not incorporate quantitative environment-specific TF regulatory influences that are modeled by EGRIN, and is therefore also limited in accurately predicting environment-specific consequences of TF perturbations. For the reasons stated above, PROM, rFBA, and IDREAM are limited in their ability to predict environment-specific phenotypic consequences of perturbations to TFs.

Additionally, there are algorithms (OptORF^18^, EMILiO^19^ and BeReTa^20^) that have the potential to predict the consequence of regulatory and metabolic network perturbations. They were originally designed to identify perturbations that maximize flux towards a desired metabolite and some of their features make them not well-suited for predicting systems-wide conditional outcomes of TF perturbation. For instance, OptORF^18^ and EMILiO^19^ use binary or fixed weights to model TF influences, which does not capture changes in relative strength of transcriptional regulation of metabolic genes across environments. By contrast, BeReTa^20^ does take into account weighted, combinatorial influences of TFs, but the analysis is restricted to genes encoding reactions of specific pathways of interest to an industrial application. Thus, none of these algorithms were designed to predict systems level phenotypic consequences (e.g., fitness and growth rate) of perturbations to the transcriptional network.

Here, we report the development of **P**henotype of **R**egulatory influences **I**ntegrated with **M**etabolism and **E**nvironment (**PRIME**), which incorporates environment-dependent combinatorial regulation of metabolic genes to mechanistically predict how individual TFs contribute to the phenotype of Mtb in any given environment. Through the use of comprehensive experimental validations, we demonstrate that PRIME significantly outperforms the previous methods in accurately predicting regulatory and metabolic genes that are conditionally required for growth on carbon sources that are specific for *in vitro* (glycerol) and *in vivo* (cholesterol) growth of Mtb. Further, PRIME has uncovered the interplay of regulatory and metabolic mechanisms that underlies Mtb’s response to drug treatment. The accuracy of PRIME in predicting quantitative phenotypic effects of TF perturbations is demonstrated by high correlation between predicted and experimentally validated consequences of knocking out all metabolism-associated TFs (one-at-a-time) on isoniazid (INH) treatment-specific fitness of Mtb strains. Through this analysis, we have discovered new vulnerabilities in Mtb that can potentiate INH action, which are supported by experimental validation.

## RESULTS

### CONDITION-SPECIFIC INTEGRATION OF REGULATION AND METABOLISM USING PRIME

A causal and mechanistic model of the transcriptional regulatory network and its quantitative influence on metabolic flux is required to characterize how the 214^21^ TFs encoded in the Mtb genome enable its physiological adaptations to disparate host relevant contexts including antibiotic treatment. We applied linear regression with TF activity (TFA) estimation using the Inferelator^15,22^ to construct an EGRIN from a compendium of 664 transcriptomes for Mtb that represented transcriptional changes in 3,902 genes (potentially regulated by 142 TFs) across 77 environmental conditions including drug treatment, pH, oxygen and carbon source utilization (**Table S1**) (http://www.colombos.net/). Relative changes in the expression of every gene across all conditions were modeled as the sum of weighted influences of a minimal set of TFs. Altogether, 142 TFs were implicated in the regulation of 3,902 genes in the genome, acting through a combinatorial scheme represented by 4,820 regulatory influences, (see **Table S2** for details). EGRIN recapitulated 2,410 of the 4,546 TF– gene interactions in the Mtb P-D network with both physical binding (from ChIP-seq experiments) and functional evidence (from transcriptional profiling)^21,23^, and added weights (β) to the influence of each TF on regulation of its target genes; here onwards we refer to this subset of 2,410 TF-gene interactions as the “EGRIN-PD Network” (**Table S2**). Thus, the Inferelator analysis added 2,410 novel TF regulatory influences that were not represented in the originally compiled P-D interaction network, accounting for 4,820 interactions in total, here onwards considered as the “EGRIN” network. Briefly, out of 4,820 interactions of EGRIN, 2410 interactions have P-D evidence (EGRIN P-D).

We investigated the degree to which EGRIN and EGRIN-PD models captured the regulation of 1,011 genes that encode enzymes implicated in catalyzing 1,229 reactions in the iEK1011^24^ model of the *Mtb* metabolic network. This analysis demonstrated that whereas EGRIN-PD modeled 1,252 regulatory influences of 104 TFs on 605 genes associated with 409 metabolic reactions, EGRIN modeled 2,568 regulatory influences of 129 TFs on 750 genes associated with 725 metabolic reactions. We leveraged the EGRIN and EGRIN-PD wiring diagrams and weights of regulatory influences inferred by the Inferelator to predict how change in the activity of a TF in a given environment manifests in altered flux through a metabolic reaction catalyzed by their regulated gene product. In order to integrate regulation with metabolism, we had to account for combinatorial regulation of metabolic genes, with each of 349 out of the 750 metabolic genes predicted to be putatively regulated by ≥2 TFs and 111 TFs predicted to regulate ≥2 metabolic genes (**Figure S1 and Table S3**), and association of ≥2 gene products to each of 313 reactions in Mtb.

The quantitative influence of a TF on the regulation of a target gene in a given environment was calculated by multiplying the EGRIN-inferred regression weight (β) of the TF influence with its absolute expression level in that environmental condition (i.e., a scaled value of signal intensity for microarray data or read counts for RNA-seq) based on distribution of values across the transcriptome compendium (**Figure 1A**; Methods). For a metabolic gene that is regulated by multiple TFs, we calculated the relative contribution of each TF to the regulation of that gene in a given environment by dividing its quantitative influence with the sum of quantitative influences of all TFs that regulate that gene. In this scheme, a TF will have a large relative consequence on the expression of a metabolic gene in an environment in which the TF is active and in high abundance, and the influences of other TFs are minimal. But the relative contribution of the TF will be proportionally lower if other TFs are also actively regulating that gene in that environment. Thus, this approach accounted for regulation of a metabolic gene by multiple TFs, and it simultaneously corrected for environment-specific changes in combinatorial regulatory schemes. For a TF that regulates multiple genes encoding enzymes or enzyme subunits for the same reaction, we considered the largest regulatory influence of that TF on any of those genes to predict its influence on flux through that reaction. Thus, together these advancements accounted for complex combinatorial associations between regulation and metabolism to assign a single relative influence factor (γ) to each TF-reaction association. The consequence of TF regulation (or knockout) on flux through a reaction is calculated by multiplying the TF-induced relative inhibition of that reaction (1-γ) to the maximum possible flux through that reaction. In this manner, by updating upper bounds of flux through all reactions catalyzed by regulated gene products of a specific TF, PRIME constrains the metabolic network to a new solution space, to enable the prediction of “environment-specific” growth consequences of perturbing a given TF which can be compared to conditional genome-wide fitness data for PRIME performance assessment (**Figure 1B**).

**Figure 1:**
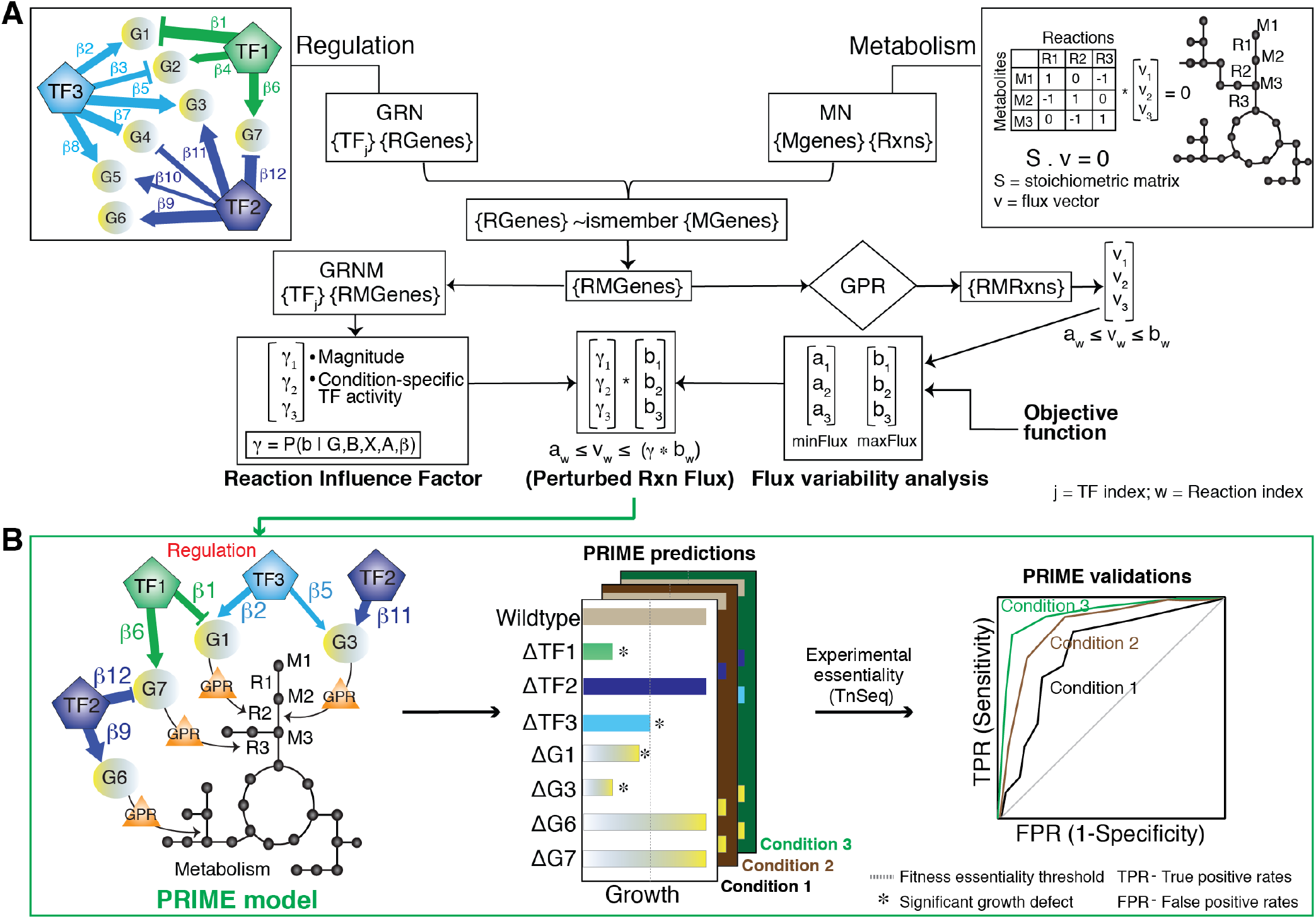
Schematic for PRIME model development and performance assessment. **A.** Schema for integration of gene regulation and metabolism. The gene regulatory network (**GRN**) models weighted regulatory influences of TFs on regulated genes (**RGenes**). A subset of the RGenes are enzyme-coding metabolic genes (**Mgenes**), whose functions are also modeled through gene-to-protein-to-reaction (GPR) mapping in a stoichiometric matrix representation of the metabolic network (**MN**). PRIME uses the integrated Gene Regulatory Network of Metabolism (**GRNM**) and a reaction flux influence estimator (**ReFInE**) to calculate the *γ* factor, which quantifies how the differential expression of multiple TFs and their weighted regulatory influences on a regulated metabolic gene (**RMGene**) manifests in altered flux (**a**: minimum flux; **b**: maximum flux) through the associated metabolic reaction (**RMRxn**) in a given environmental condition. **B.** Illustration of condition-specific gene phenotype predictions and performance assessment. The example illustrates how PRIME predicts relative growth consequence of single gene knockouts in TFs (e.g., TF1, TF2 and TF3) and RMGenes (e.g., G1, G3, G6 and G7) in different contexts (e.g., Condition 1, 2, and 3). The vertical line in the barplot depicts a user-defined threshold in growth inhibition, below which a gene is deemed essential. Performance of PRIME is quantified using a Receiver Operating Characteristic (**ROC**) curve based on accuracy of PRIME-predicted essential and non-essential genes in a given condition to experimentally determined phenotype consequences using transposon mutagenesis coupled with sequencing (**TnSeq**) in the same condition.

### PERFORMANCE ASSESSMENT OF PRIME

In order to compare performance of PRIME to previously developed methods, we had to first update the PROM model with the latest version of the Mtb P-D interaction map^12,21^ and the current version of the metabolic network model iEK1011^24^ (1,011 genes encoding enzymes for 1,229 reactions) that was used to construct PRIME. Using the methodology described in the original PROM paper^11,12^, 2,416 out of 2,555 P-D interactions for 104 TFs were mapped to 605 genes assigned to 632 reactions in the iEK1011 metabolic network model. This represents a significant improvement in the overall coverage of TFs and metabolic genes in the PROM model (**Table 1**, **Figure 2A**). In parallel, we also developed the first IDREAM model for Mtb by incorporating confidence scores for 2,407 regulatory influences for 142 TFs within the EGRIN model (FDR <0.25) on a total of 641 genes associated with 639 reactions within iEK1011 (**Table 1, Figure 2B**). The slightly higher numbers of TFs and metabolic genes in IDREAM and PRIME (**Figure 2B**) are because they use the EGRIN model, which has better coverage of genomewide TF regulation across diverse environments, relative to the P-D interaction map generated in standard growth conditions that was used in PROM (**Figure 2C**). In summary, the updated PROM and IDREAM models were similar to PRIME in terms of coverage of the total number of TFs and metabolic genes and suitable for comparing performance across the models. (**Table 1**).

**Figure 2:**
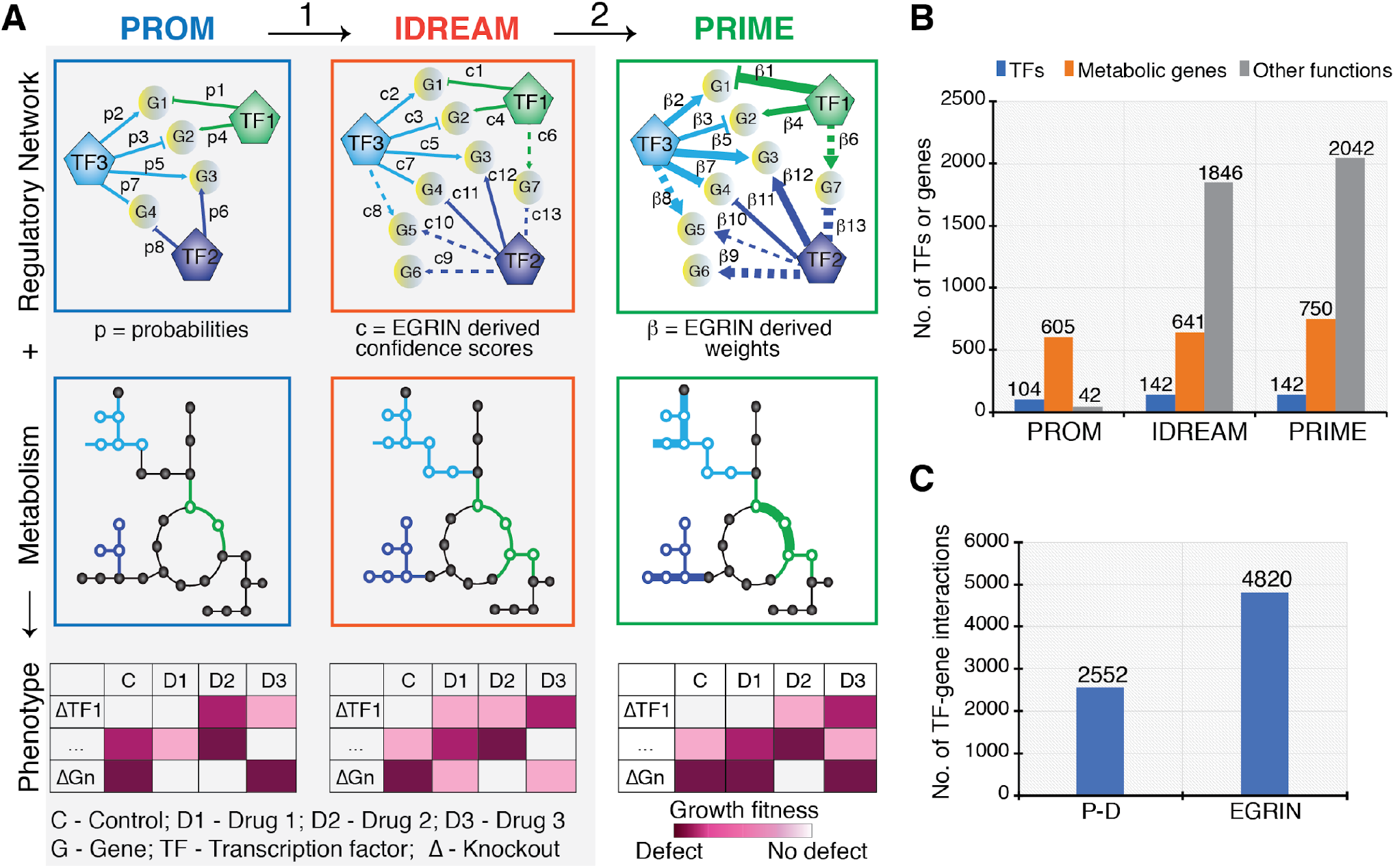
PRIME model advancements. **A.** Advancements in PRIME over previous methods (PROM and IDREAM) are indicated as (1) incorporation of regulatory influences from EGRIN (regression-based interactions are shown as dotted lines), which increases coverage of the regulatory network, (2) incorporation of the magnitude of regulatory influence of TFs on metabolic genes (β – shown as varying edge thickness) instead of probability (p) and confidence score (c) significantly improved the predictive accuracy of environment-specific gene essentiality. **B.** Number of TFs and genes from PRIME, IDREAM and PROM. **C**. Number of TF-gene interactions identified using regression-based EGRIN and Protein-DNA (P-D) interactions from ChIP-seq data.

**Table 1:** Summary of PROM, IDREAM, and PRIME model features

| Mtb Model Features | Chandrasekaran,<br>2010 | Ma, 2015 | Present Study |  |  |
| --- | --- | --- | --- | --- | --- |
|  | MTBPROM1.0 | MTBPROM2.0 | PROM* | IDREAM* | PRIME |
| Metabolic model | iNJ661 | iSM810 | iEK1011 | iEK1011 | iEK1011 |
| Number of reactions | 1025 | 938 | 1229 | 1229 | 1229 |
| Number of metabolic genes in the<br>metabolic network | 661 | 810<br>(759 genes in<br>iEK1011) | 1011 | 1011 | 1011 |
| Regulatory network | Balazsi 2008 | Minch 2015 | Minch 2015 | EGRIN<br>(FDR<0.25) | EGRIN<br>(Precision=<br>50%) |
| Number of transcription factors | 30 | 104 | 104 | 142<sup>#</sup> | 142<sup>#</sup> |
| Number of interactions | 218 | 2555 | 2555 | 3643<sup>#</sup> | 4820<sup>#</sup> |
| Number of genes in the regulatory<br>network (**metabolic / total**) | 178 / 178 | 647 / 647 | 605 / 647 | 641 / 2487 | 750 / 2905 |

\*The PROM model was updated in this study by incorporating the latest metabolic network (MN) model for Mtb; the IDREAM model was constructed in this study to evaluate performance relative to the other methods
<sup>#</sup>PRIME uses the same EGRIN network as IDREAM, but incorporates the weights of regulation of each metabolic enzyme to update the constraint on reaction fluxes through the MN.

We compared the performance of PRIME to PROM and IDREAM by assessing their accuracy (sensitivity and specificity) in predicting environment-specific growth inhibition upon TF deletion for Mtb cultured in minimal medium with glycerol or cholesterol as the carbon source. While Mtb is typically grown with glycerol during *in vitro* culture, the pathogen is capable of utilizing host-derived lipids, such as cholesterol, during infection. It is known that distinct metabolic genes and networks are associated with these two modes of growth. Accuracy of model predictions were evaluated using a leave-one-out cross validation (LOOCV) strategy^25^ for comparison of model predictions to experimentally determined phenotypic consequences of transposon mutagenesis in genome-wide fitness screens (TnSeq) of Mtb cultured with glycerol or cholesterol^26,27^. Specifically, for each model we generated a set of receiver-operating characteristic (ROC) curves by plotting the true positive rate (i.e., proportion of model-predicted essential genes that were verified by experiment) and false positive rate (i.e., proportion of model-predicted essential genes that were experimentally determined to be non-essential) by leaving out one TF in each analysis. The distribution of area under the ROC curves (ROC-AUC) from the LOOCV analysis of model predictions of which TFs are essential for Mtb growth on cholesterol was used as a metric of performance. First, we evaluated predictions from PRIME using either EGRIN-PD or EGRIN, inferred using different Inferelator parameter settings as the source of regulatory influences, and concluded that the latter contributed to significantly better performance (**Figure S2**). Therefore, here onwards all results reported for PRIME are based on regulatory influences from the EGRIN network. The LOOCV analysis demonstrated that the performance of PRIME was significantly better relative to PROM and IDREAM in both cholesterol and glycerol carbon sources (**Figure 3A, 3B** and **Figure S3**). In addition to providing a rigorous means for performance evaluation, the LOOCV^25^ analysis also identified a clear division of TFs in terms of their ROC-AUC values for the PRIME model. Further analysis revealed that the top performing TFs (20 and 12 TFs for glycerol and cholesterol, respectively) contributed maximally (up to 65% of overall biomass accumulation) to the overall fitness of Mtb (**Table S4**). Out of 119 TFs with TnSeq data, the cholesterol fitness of 65% (77 TF KOs) were accurately predicted by PRIME, whereas IDREAM and PROM accurately predicted only 45% (53 TFs) and 30% (35 TFs), respectively (**Figure 3C**). Similarly, PRIME accurately predicted glycerol fitness for 92 out of 119 TFs (77%), whereas IDREAM accurately predicted 55% (65 TFs) and PROM predicted 36% (43 TFs) (**Figure 3D**). In general, PRIME, IDREAM and PROM predictions differed significantly (p-value <2.2e-16, t-test) both in the numbers and the context in which genes were called essential or non-essential.

**Figure 3:**
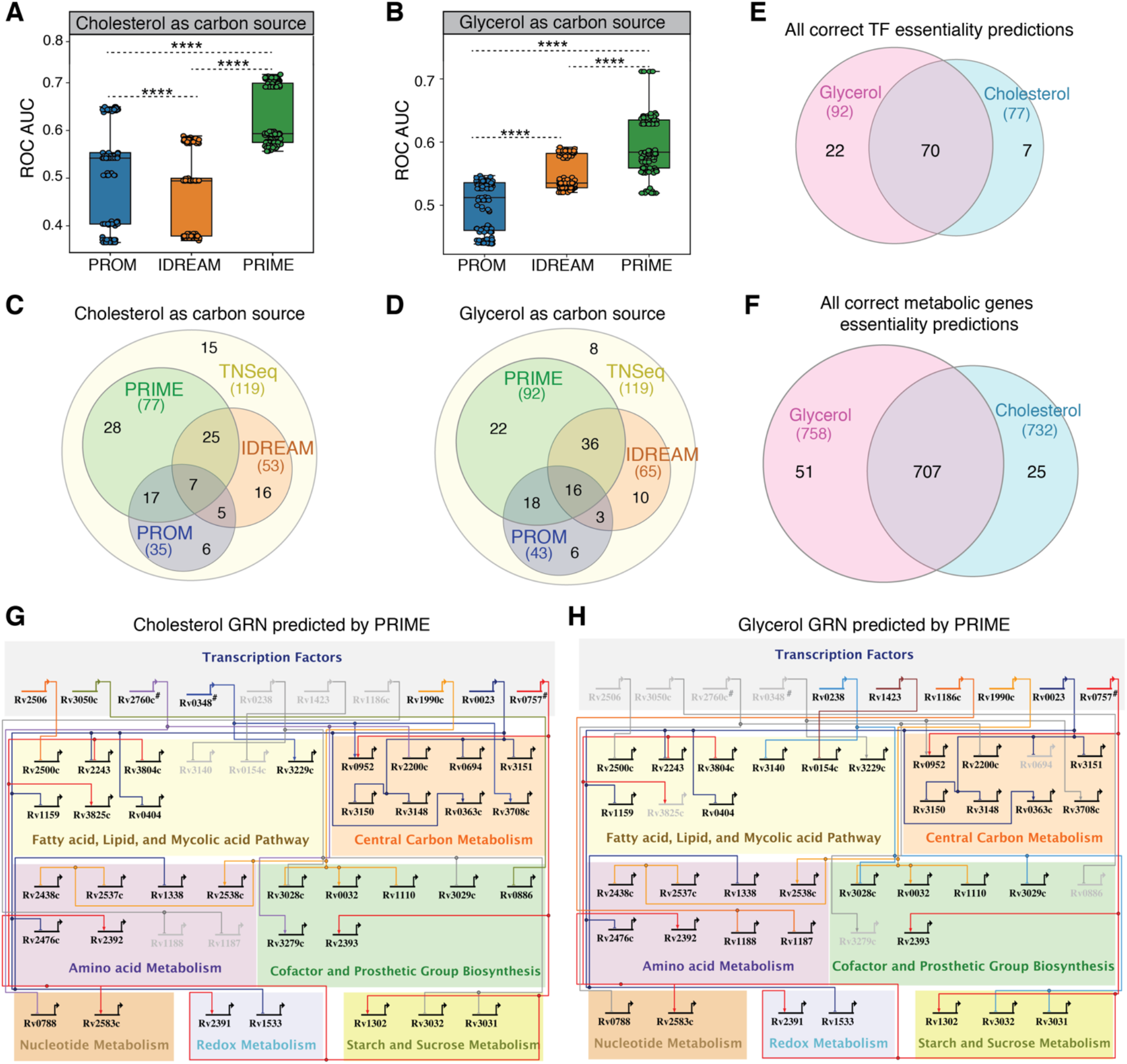
Validation of PRIME predictions of conditional gene essentiality. Sensitivity and specificity of PRIME, PROM, and IDREAM predicted TF essentiality in **A**. cholesterol and **B.** glycerol as determined by LOOCV analysis for the area under the receiver operating characteristic curve (ROC AUC). Statistical significance was calculated as *p*-value with two sample t-test. ****: *p*-value < 0.0001. Comparison of all positive predictions (true positives and true negatives) for TF essentiality by PRIME, PROM, and IDREAM in **C.** cholesterol and **D.** glycerol. **E.** The number of all correct PRIME predictions (true positives and true negatives) of TF knockouts across the two conditions (glycerol and cholesterol) that are validated by experimental TnSeq data. **F**. The number of all correct PRIME predictions for deletion of all genes in the metabolic network across the two conditions that are validated by experimental TnSeq data. BioTapestry visualization showing a subset of the gene regulatory network of Mtb under growth in **G.** cholesterol and **H.** glycerol. TFs are grouped together in the top panel (represented by bent arrows), which extend to horizontal and vertical lines that connect to their regulatory gene targets. Highlighted TFs were predicted by the PRIME model to be essential and validated through TnSeq dataset in relevant conditions.

Using PRIME, 22 and 7 TFs were accurately predicted (either essential or non-essential) for growth only with either glycerol or cholesterol, respectively, as determined by experimental fitness screening (**Figure 3E**). Similarly, 51 and 25 metabolic genes were accurately predicted by PRIME for growth on either glycerol or cholesterol, respectively (**Figure 3F**). Among the PRIME predicted essential TFs, Rv2506, Rv3050c, Rv2760c, and Rv0348 are essential for growth on cholesterol, presumably because they conditionally regulate genes encoding enzymes or enzyme subunits catalyzing essential metabolic processes during cholesterol utilization (**Figure 3G**). For example, Rv2506 represses genes likely to be involved in branched-chain amino acid catabolism, which leads to the production of acetyl-coA and propionyl-coA^28^. Propionyl-coA is also an endpoint of cholesterol degradation and can be toxic to Mtb^29^. It is possible that Rv2506 repression of branched-chain amino acid metabolism genes prevents accumulation of toxic metabolic intermediates during growth on cholesterol. All in all, perturbation of cholesterol utilization in Mtb could induce metabolite intoxication^29^, unbalanced central metabolism^30^ or lead to carbon starvation^31^. As such, TFs such as Rv2506, Rv3050c, Rv2760c and Rv0348 represent potential vulnerabilities in the cholesterol utilization pathways of Mtb that could be targeted by drugs. Notably, these TFs were also ascertained to be essential by the TnSeq screen performed with cholesterol as the carbon source^26^ and are non-essential in glycerol (shown as inactive nodes in **Figure 3H**). Other TFs (Rv1990c, Rv0023 and Rv0757) were predicted (and validated by TnSeq^26^) to be essential for growth with both carbon sources or only essential for growth on glycerol (e.g., Rv0238 and Rv1423).

### PRIME RANK IDENTIFIES THE ESSENTIAL TRANSCRIPTIONAL FACTORS AND GENES FOR SURVIVAL DURING DRUG TREATMENT

We used PRIME to investigate the regulatory and metabolic networks that drive physiological adjustments (e.g., cell wall modifications, shifts in metabolism and respiration) to enable the pathogen to survive and persist during drug treatment. To expose novel network vulnerabilities of Mtb in response to drug treatment, we generated transcriptome profiles of Mtb treated for 24 h with high- and low-doses of seven drugs (**Table S5**). The transcriptome profiles were analyzed using the PRIME model to identify the metabolic networks and their associated regulators that were essential for growth in the absence and presence of drug treatment. This analysis found clear distinction in TF essentiality between the untreated and drug-treated PRIME models and revealed that drug doses largely grouped together (**Figure 4A**). Interestingly, the TF essentiality profiles of rifampicin (a transcription inhibitor) were dosedependent; the rifampicin profile at low-dose clustered separately, while the high-dose profile clustered with linezolid (a protein synthesis inhibitor). The resemblance to linezolid at high-dose suggests that a secondary effect of strong rifampicin-induced transcription inhibition also impacts translation. Furthermore, we observed that the TF essentiality profiles of isoniazid (inhibitor of cell wall synthesis) were quite distinct to the other six drugs. In fact, 58 TFs become conditionally essential in the presence of isoniazid because of their mechanistic role in regulating 569 metabolic reactions required for supporting growth during isoniazid treatment. This highlights the multitude of regulatory-metabolic networks associated with cell wall disruption in Mtb and the extreme vulnerability in cell wall metabolism.

**Figure 4.**
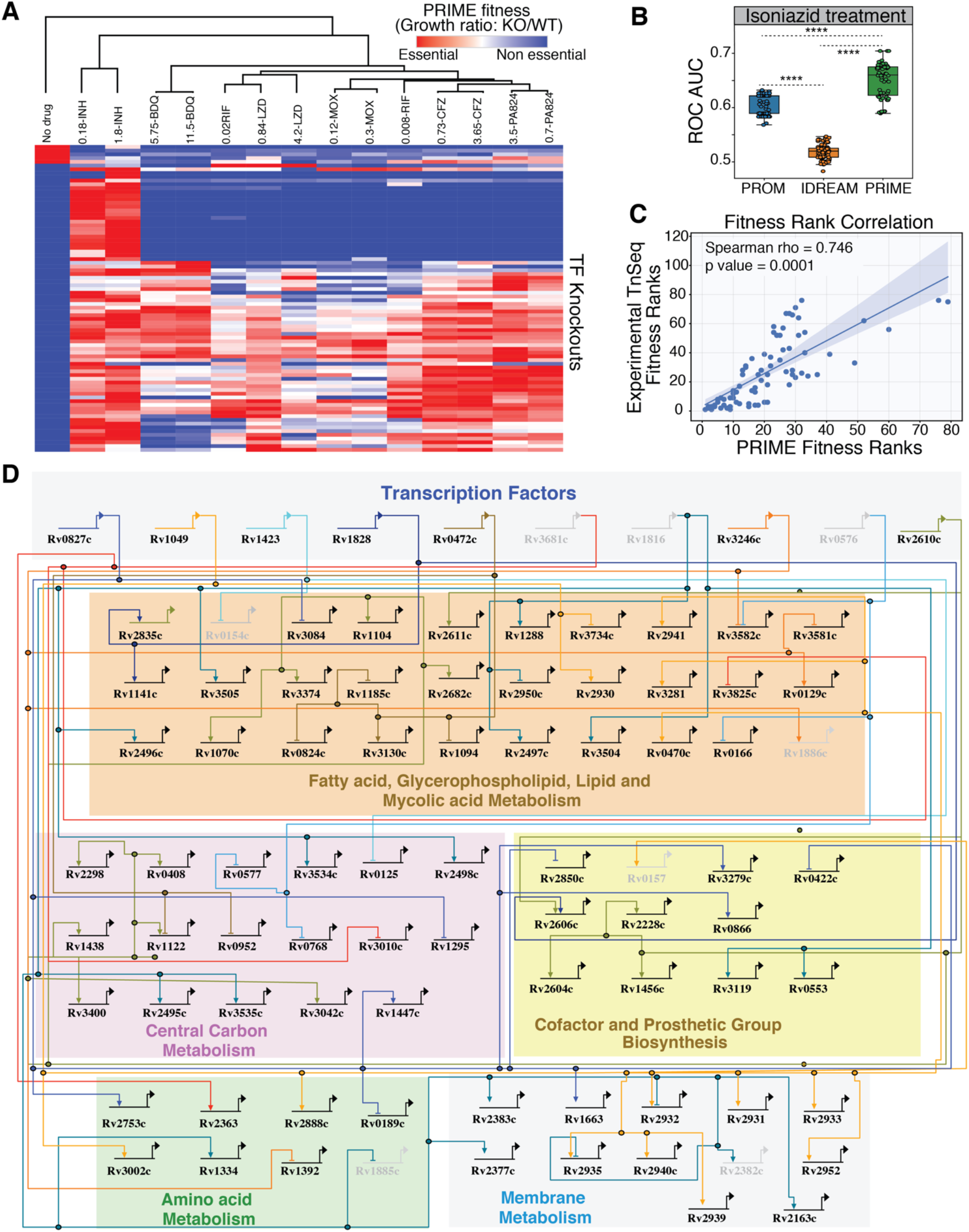
Drug-specific predictions of PRIME. **A.** Heatmap of PRIME derived fitness for all TF knockouts in the presence of 7 primary drugs and control at 24 h. The numbers indicate the concentration of drug used in μg/mL. INH: isoniazid, BDQ: bedaquiline, RIF: rifampicin, LZD: linezolid, MOX: moxifloxacin, CFZ: clofazamine, PA824: pretomanid. **B.** Sensitivity and specificity of PRIME, PROM, and IDREAM predicted TF essentiality in the presence of INH as determined by LOOCV analysis for the area under the receiver operating characteristic curve (ROC AUC). Statistical significance was calculated as *p*-value with two-sample t-test. ****: *p*-value < 0.0001. **C.** Correlation of TnSeq experimental fitness ranking of TFs and PRIME derived fitness ranks. **D.** BioTapestry visualization showing a subset of the gene regulatory network of Mtb with PRIME predictions during INH treatment. Some of the highlighted TFs were predicted as essential in the presence of INH (Rv0827c, Rv1049 and Rv0472c), while others were predicted essential in both the absence and presence of INH (Rv1423, Rv1828, Rv3246c, and Rv2610c). The lightened TFs were predicted essential in the untreated control but non-essential in the presence of INH (Rv3681c, Rv1816, and Rv0576). All of these PRIME predictions were validated by experimental fitness screening in relevant conditions.

Focusing on isoniazid (INH), we evaluated the accuracy of these predictions against experimentally-determined fitness values from a genome-wide TnSeq screen performed in the presence of a subinhibitory concentration of INH^32^. Notwithstanding the difference in dosage of drug treatment of the input transcriptome data used in the PRIME model (0.18 ug/mL, 1.8 ug/mL) and in the TnSeq fitness screen (27 ng/mL), the LOOCV analysis demonstrated high sensitivity and specificity of PRIME predictions of gene essentiality (max ROC AUC = 0.685), significantly outperforming PROM (max ROC AUC = 0.625) and IDREAM (ROC AUC = 0.6) (**Figure 4B)**. We also used PRIME to rank order TFs based on their relative importance in supporting growth in the presence of INH, and compared these ranks to TnSeq determined importance of TFs. There was striking correlation (Spearman’s rho = 0.695; p-value = 0.0001) in the rank ordering of TFs based on the predicted (PRIME) and observed (TnSeq) magnitude of growth inhibition of Mtb in the presence of INH upon knocking out each TF one-at-a-time (**Figure S4**). The correlation increased dramatically (Spearman’s rho = 0.746, p-value = 0.0001) when only TFs implicated by EGRIN as regulators of essential metabolic reactions were considered in this analysis, demonstrating the remarkable accuracy of PRIME in capturing how the differential regulation by TFs modulates flux through essential metabolic reactions to manifest at a phenotypic level (**Figure 4C**). Notably, PRIME accurately predicted that knocking out the top 10 TFs one-at-a-time would result in at least 65% and up to 95% Mtb growth inhibition during INH treatment, but not in the absence of drug treatment, implicating these as conditional vulnerabilities for significantly potentiating INH treatment (**Table S6**).

To aid in the interpretation of PRIME predictions, we developed the PRIME pathway analysis (PPA) tool to uncover in a single-step the specific metabolic reaction(s) regulated by a TF that make it essential for growth in a given environmental condition. Given a TF, PPA identifies all reactions catalyzed by the genes it is predicted to regulate, rank orders the target genes based on the relative contribution of their gene product in driving flux towards biomass accumulation, and outputs a TF-metabolic gene-reaction map as a putative mechanism by which the TF is likely to be essential in a given environmental context. Using PPA, we identified the specific metabolic reactions that were mechanistically responsible for the conditional essentiality of 23 TFs validated by TnSeq data^32^ to be essential in the presence of INH. For example, we discovered the mechanisms underlying the essentiality of Rv0827c, Rv1049, Rv1423, Rv1828 and Rv0472c for growth in the presence of INH (**Figure 4D**). Altogether, PPA uncovered that 58 of the 142 TFs were conditionally essential for growth on INH because they conditionally regulate 569 key reactions across 55 pathways, including 84 reactions within fatty acid metabolism and mycolic acid biosynthesis (target of INH). In so doing, PRIME has provided the most comprehensive systems level perspective into strategies to potentiate INH killing by targeting TFs that mediate Mtb’s metabolic response to INH treatment.

## DISCUSSION

We have demonstrated that by incorporating how TFs act contextually in combinatorial schemes to regulate gene expression, PRIME outperformed PROM and IDREAM in accurately predicting how transcriptional regulation redirects metabolic flux to manifest in environment-specific phenotypes of Mtb. The shortcoming of PROM can be attributed to its reliance on P-D interactions for regulatory network, which are plagued with false positive interactions (because overexpression of TFs can force nonfunctional binding across the genome) and false negative interactions because of lack of appropriate context (e.g., missing co-factors). Hence, a P-D interaction does not capture whether a TF is regulating a gene in a given condition, which is better modeled by regulatory influences inferred using regression analysis of transcript level changes in TFs and all genes across the genome. However, despite incorporating regulatory influences from the same EGRIN network, IDREAM performance was inferior compared to PRIME, and in fact its performance in predicting gene essentiality in cholesterol and INH was worse than PROM. One explanation could be that relative to the number of P-D interactions used in PROM, IDREAM used nearly twice as many EGRIN-based regulatory influences that were inferred from a wide range of environmental contexts, without taking into account combinatorial regulatory schemes, weights of regulatory influences, or the absolute expression levels of TFs to prune regulatory edges that were not relevant for a given environmental context. Hence, reliance on a P-D interaction map, and even just the likelihood that a TF might regulate a gene based on regression analysis are both insufficient to capture the complex environment-dependent interplay of transcription and metabolism. Altogether, these comparative analyses have demonstrated that four key advancements in PRIME addressed the shortcomings of PROM and IDREAM: (i) PRIME took full advantage of EGRIN predictions to incorporate weights of TF regulatory influence on each gene; (ii) PRIME calibrated the relative influence of each TF on a given metabolic gene by accounting for all TFs that were also implicated in the regulation of that gene; (iii) PRIME accounted for regulation of multiple genes that encode enzymes for the same reaction by considering which gene(s) contributed maximally towards flux through that reaction in a given environmental context; and, finally (iv) PRIME considered the absolute expression level of each TF to evaluate the degree to which each regulatory influence was active in a given environment.

By demonstrating better accuracy in predicting environment-specific phenotypes of Mtb using EGRIN, PRIME overrides the need for a physical map of P-D interactions, which is difficult to generate for many organisms, across all environments of interest, and especially in some contexts, such as within infected tissue. In fact, the incompleteness of the P-D interaction map was demonstrated by the significant drop in the performance of PRIME upon excluding regulatory influences that were not supported by physical TF-gene interactions (i.e., EGRIN P-D). By contrast, EGRIN is inferred directly from a compendium of transcriptomes, which can be profiled across relevant environmental conditions with minimal manipulation (e.g., without overexpression of TFs) and even within infected cells using technologies like Path-seq^33^. As a consequence, EGRIN discovers a significantly larger number of novel regulatory mechanisms, including the combinatorial schemes and specific environmental contexts in which they are conditionally active. This explains why PRIME discovered mechanisms that become conditionally essential in the presence of INH, but also accurately predicted the relative importance of each TF for enhancing the potency of INH. Based on this observation, we posit that PRIME will be especially valuable to prioritize genes that represent novel context-dependent vulnerabilities that could be targeted to potentiate the action of any antibiotic and achieve faster clearance with a lower dosage. By enabling the *in-silico* discovery of vulnerabilities within the Mtb network, PRIME also overrides the need for large scale transposon mutagenesis-based experiments (e.g., TnSeq, TraSH, HITS, etc), which are resource-intensive and difficult to perform across all conditions relevant to the lifecycle of Mtb. Instead, PRIME can be used to rank prioritize the strains and contexts in which to assay for an expected phenotype. This capability is particularly powerful considering the numerous mechanisms by which Mtb can be phenotypically different, with different antibiotic sensitivities. Additionally, there is growing evidence that upon gaining resistance to an antibiotic, the regulatory and metabolic networks within a pathogen are remodeled in order to reallocate resources for supporting the new phenotype^34^. Using PRIME, we can delineate novel vulnerabilities within these remodeled regulatory and metabolic networks to devise strategies for rationally disrupting the antibiotic resistance phenotype with a second drug.

PRIME will also be useful in biotechnology applications to further optimize the production of desired end products by rewiring the regulatory networks of metabolically engineered strains. Advancements in metabolic engineering have been effective in substantially increasing flux towards the production of a desired metabolite^18–20,35^ but there is a limit to which metabolic engineering alone can improve the overall yield. It has been proposed that further enhancements in yield would require reprogramming of the regulatory network to control when genes of the engineered pathways are expressed, and to rationally up and down regulate competing metabolic pathways to maximize flux and resource allocation towards the desired objective. Hence, by using PRIME, metabolic engineering of high-yielding strain phenotypes can be identified. Although the capabilities of PRIME are elucidated extensively using Mtb as a model system in this study, we foresee the use and applications of PRIME in various organisms due to its scalability.

## METHODS

### CONSTRUCTION OF EGRIN GENE REGULATORY NETWORK FOR *MYCOBACTERIUM TUBERCULOSIS*

The Mtb EGRIN used in this study was constructed using the Inferelator algorithm^15,22^ trained on a transcriptional compendium for Mtb with 3,902 genes across 664 experimental conditions (downloaded from the COLOMBOS database) and an experimentally supported signed Mtb P-D network (generated as previously described in ^33^). The original transcriptional compendium contained a larger number of genes and conditions but was modified to remove genes and conditions with missing values. Briefly, we used the Inferelator to identify potential transcriptional regulators for the 3,902 Mtb genes in the expression compendium, as previously performed for other species^22,36^. The Inferelator first estimates the regulatory activities of each transcription factor activity (TFA) using the expression profile of TF known targets (encoded in the signed P-D network). Then, the Inferelator uses a Bayesian Best Subset Regression to estimate the magnitude and sign (activation or repression) of potential interactions between TFs and genes. As before, we bootstrapped the expression data (20 times) to avoid regression overfitting. The Inferelator generates two scores for each TF-gene interaction, the corresponding regression coefficient (weight – β) and a confidence score. The second score indicates the likelihood of the interaction. The final set of TF-gene interactions was defined with a 0.5 precision cutoff. This means that 50% of all interactions in the inferred network were already present in the signed P-D network used for training, while the other half corresponded to putative novel TF-gene interactions.

### DEVELOPMENT OF PRIME

The PRIME algorithm has been developed by integrating weights (β) from EGRIN with metabolic network (MN) models for phenotype prediction in a context-specific manner (wiring diagram in Fig. 1). PRIME requires 1) a MN in the format of constraints-based model^37,38^ in systems biology markup language (SBML), an XML format as input, that are represented *in silico* in the form of a stoichiometric matrix, wherein every column corresponds to a reaction and every row corresponds to a metabolite. These constraints-based models were used to integrate the regulatory influences by updating the reaction flux, 2) a regulatory network containing TF and gene interactions (one array of regulators and one array of corresponding gene targets), 3) magnitude/weights (β) of regulatory influences for each of the interactions (array of magnitudes) derived from Inferelator and 4) the gene expression data profiled under a specific condition (gene ids and their expression, provided as ratio to the control – in case of environment-specific predictions the ratio between initial t0 and final time point tn). The pipeline of PRIME initially links each metabolic gene in MN to its associated regulators considering the combinatorial effects, followed by applying the calculated relative influence factor. Specifically, we have introduced a new way to calculate the relative influence factor (*γ*), a value that quantitatively constrains the reaction flux constraint space. The equations 1 to 5 consists of the details involved in each successive step within the algorithm.

Given a TF *j* influencing a metabolic gene *i* of reaction *w*, we define *γ_i,w_* as,

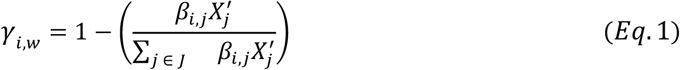

where *J* is the subset of TFs that influence gene *i*, and 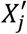 is the scaled expression of a TF *j* in a particular condition *c* of a coherent environmental context *B* as,

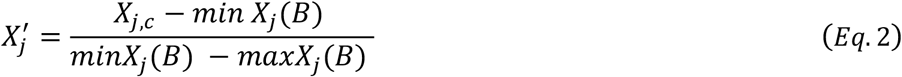

Then, the regulatory influence that exerts the larger effect on reaction *w* across the set of metabolic genes *i*, ∈ *I* of a given reaction has been identified as,

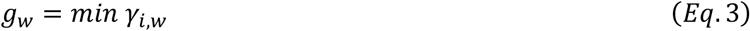

At this point, it is straightforward to incorporate calculated weights as new upper bounds,

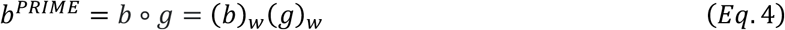

to the flux balance analysis (FBA)^38^ formalism, assuming steady state metabolic concentrations, and defining the system mass balance as *S.v* = 0, to maximize the objective function *Z* = *c^T^v* such that fluxes are within the new boundary conditions,

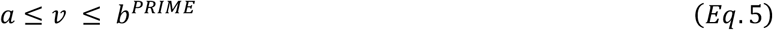

The objectives in each prediction are defined during FBA optimization. The phenotype predictions mentioned in this study are the optimized biomass predicted by FBA. The complete PRIME algorithm package and details of the required input dataset is available for download from our GitHub Repository (https://github.com/baliga-lab/PRIME). All model simulations related to FBA were performed on MATLAB_R2019a platform using the recent version of COBRA^39^ (The COnstraint-Based Reconstruction and Analysis) toolbox. *In silico* gene essentiality predictions were performed using the COBRA toolbox ‘single-gene-deletion’ function in MATLAB.

### INCORPORATING DRUG TREATMENT GENE EXPRESSION DATA ON METABOLIC MODEL

The iEK1011^24^ metabolic network (MN) model was used for all the predictions in this study. For drugspecific models, we applied the gene expression data from both drug-treated and untreated control experiments using the GIMME^40^ algorithm on the iEK1011 MN model. This step was carried out to constrain the MN model to the specific condition being tested. We used GIMME because of the flexibility in defining objective function during implementation. The GIMME algorithm is implemented in the MATLAB_R2019a platform, using the “GIMME.m function” in the COBRA Toolbox after processing the gene expression data through ‘mapExpressionToReactions.m’ function to convert the gene expression values as inputs to GIMME.

### PROM MODELS

For developing PROM^11,12^ models, we followed the PROM approach^11^ to estimate the probability that a target gene is ‘ON’ or ‘OFF’ in the absence of the TF i.e., in the event of a TF knockout. This was calculated from a gene expression dataset as, Probability, P (Gene = 1|TF = 0) or P (TF = 1|Gene = 0). The gene expression threshold that delineated between the ‘ON’ and ‘OFF’ states was set as quantile (0.33) from the input expression data. These probabilities were then used to constrain the maximal fluxes of the reactions catalyzed by the gene products in the metabolic model as *p × Vmax,* where p is the probability of the gene being on. The user defined “kappa” value was used as similar to earlier PROM models^11^. All PROM predictions and simulations were performed using PROM.m (MATLAB script) on the MATLAB_R2019a platform. We used iEK1011 metabolic network model in XML format as input in the PROM. The P-D derived regulatory network was obtained from the study^21^, similar to the MTBPROMV2.0^12^.

### IDREAM MODELS

For IDREAM^13^ models, the GRN derived using EGRIN, was integrated with the PROM pipeline as it had been done previously for the yeast system^13^. We ran 200 iterations in EGRIN to calculate the confidence score for all predictions. For each gene, we estimated a false discovery rate (FDR) for each TF by counting the fraction of models that identified that factor as a regulator. Thus, if TF1 was predicted to regulate gene1 in 191 of 200 models, then the TF-gene interaction identified would have an FDR = 0.045. We included only those interactions that passed an FDR cutoff of 0.25. We used EGRIN-derived GRN to integrate it with iEK1011 metabolic network model of Mtb using the PROM framework. The user defined “kappa” value was used as similar to earlier PROM models^11^. IDREAM does not rely on probabilities, hence the gene expression dataset was not used in IDREAM instead ‘prob_prior’ in the PROM function was set based on the EGRIN FDR values for each TF-gene interaction. If the TF is an activator of a gene, we use the FDR value directly, if it is an inhibitor, we use 1 -FDR value as ‘prob_prior’. EGRIN network was derived using Inferelator in R (Inferelator.pkg.R) and PROM predictions and simulations were performed using PROM.m (MATLAB script) on the MATLAB_R2019a platform as similar to PROM model development.

### PERFORMANCE ASSESSMENT OF PRIME PREDICTIONS

The predictive power of PRIME as a binary classifier (essential or non-essential) between the model predicted gene essentiality and experimentally defined gene essentiality (TnSeq) has been performed using receiver operating characteristic (ROC) curve. A gene was considered “essential” if its deletion reduced the biomass by >85%. By this analysis, the model classified each gene as “essential” or “nonessential”. We compared the gene essentiality predictions from Mtb grown under glycerol and cholesterol as carbon source with the available experimental TnSeq data^26^ and deduced the confusion matrix to derive true positive rates (TPR) and false positive rates (FPR). We also took advantage of the follow-up study where Bayesian analysis was used to assign calls as essential and non-essential for the same TnSeq dataset^27^. We expanded the analysis of TnSeq data to classify essential and non-essential with a cutoff value of using cholesterol/glycerol ratio of 0.6 in order to assign calls for all the genes. This classification led to the elucidation of sensitivity and specificity of the model using ROC curve analysis.

Briefly, the gene expression data of Mtb profiled under growth on Glycerol (GSE52020) and Cholesterol (GSE13978) were used to generate condition-specific metabolic networks using GIMME. PRIME was applied on these models to predict gene and TF essentialities according to the condition tested. These predictions were then compared to the TnSeq data. A similar sensitivity and specificity analysis was performed while validating the performance of PRIME for INH-specific predictions using experimentally derived TnSeq data^32^. To construct the INH-specific metabolic models, we used INH-treated Mtb transcriptome sequencing (RNA-seq) data generated in this study (see below).

### PRIME PATHWAY ANALYSIS (PPA) PIPELINE

The PRIME pathway analysis (PPA) pipeline was developed to derive the metabolic association of a specified TF in a simple process by accessing PRIME model genes and their interactions. The top ranked TFs and their associated metabolic genes are further linked to their metabolic processes using the PPA pipeline. PPA is provided as PRIMEanalysis.m (MATLAB script). All analyses related to PPA were performed in MATLAB_R2019a platform. The illustration of PPA-derived essential gene regulatory-metabolic networks were deduced using BioTapestry tool (http://www.biotapestry.org/).

### DRUG TREATMENT CULTURING CONDITIONS

Experiments were performed using *Mycobacterium tuberculosis* H37Rv grown with mild agitation at 37°C in standard 7H9-rich media consisting of Middlebrook 7H9 broth supplemented with 10% Middlebrook ADC, 0.05% Tween-80, and 0.2% glycerol. Frozen 1 mL stocks of Mtb cells were added to 7H9-rich medium and grown until the culture reached an OD_600_ of ~0.4-0.8. The cells were then diluted to OD_600_ of 0.05 and added to 7H9-rich medium containing drugs at the predetermined amounts. Samples, in biological triplicate, were collected at 24 h after drug treatment by centrifugation at high speed for 5 min, discarding supernatant and immediately flash freezing the cell pellet in liquid nitrogen. Cell pellets were stored at −80° C until RNA extraction was performed as previously described^41^.

### PROCESSING AND ANALYSIS OF RNA-SEQ DATA

Sample collection and RNA-extraction was performed as described above. Total RNA samples were depleted of ribosomal RNA using the Ribo-Zero Bacteria rRNA Removal Kit (Illumina, San Diego, CA). Quality and purity of mRNA samples was determined with 2100 Bioanalyzer (Agilent, Santa Clara, CA). Samples were prepared with TrueSeq Stranded mRNA HT library preparation kit (Illumina, San Diego, CA). All samples were sequenced on the NextSeq sequencing instrument in a high output 150 v2 flow cell. Paired-end 75 bp reads were checked for technical artifacts using Illumina default quality filtering steps. Raw FASTQ read data were processed using the R package DuffyNGS^42^. Briefly, raw reads were passed through a 2-stage alignment pipeline: (i) a pre-alignment stage to filter out unwanted transcripts, such as rRNA; and (ii) a main genomic alignment stage against the genome of interest. Reads were aligned to *M. tuberculosis H37Rv* (ASM19595v2) with Bowtie2^43^, using the command line option “very-sensitive.” BAM files from stage (ii) were converted into read depth wiggle tracks that recorded both uniquely mapped and multiply mapped reads to each of the forward and reverse strands of the genome(s) at single-nucleotide resolution. Gene transcript abundance was then measured by summing total reads landing inside annotated gene boundaries, expressed as both RPKM and raw read counts. We used the raw read counts as input for DESeq2^44^ to obtain DESeq2 normalized counts. The RNA-seq data of Mtb response to drug exposure generated for this study are publicly available at the Gene Expression Omnibus under accession number GSE165673.

## Acknowledgements

We thank members of the Baliga lab for critical discussions and feedback. This work is funded by Bill and Melinda Gates Foundation (INV-009322) and the National Institute of Allergy and Infectious Diseases of the National Institutes of Health (R01AI128215 and U19AI135976).

## Competing interest

The authors declare no competing interest.

## Author Contributions

**NSB** and **SRCI** conceptualized the study and designed the research. **SRCI** developed the PRIME algorithm, performed all the computational analyses related to PRIME and metabolic modelling, analyzed all data represented and designed all figures. **MLAO** performed computational analyses related to regulatory networks and contributed in the initial process of PRIME conceptualization. **RR** and **MP** performed the experiments related to drug treatment and transcriptomic profiling. **ALGL** contributed in PRIME method conceptualization in early stages. **EJRP** designed and analyzed transcriptome studies, and analyzed fitness data of PRIME analysis. **NSB** and **EJRP** provided overall supervision. **SRCI**, **EJRP**, and **NSB** wrote the manuscript. All authors read the manuscript and approved its content.

## Data availability

Input files for PRIME used in this study are provided as **File S1**. All PRIME-generated data are provided as supplementary materials. PRIME code, with data and description for implementation, is available in GitHub repository: https://github.com/baliga-lab/PRIME. The RNA-seq data generated for this study are available in the Gene Expression Omnibus under accession no. GSE165673.

## Notes

https://github.com/baliga-lab/PRIME

### Competing Interest Statement

The authors have declared no competing interest.

