## Supplementary Figures for "Quantitative prediction of conditional vulnerabilities in regulatory and metabolic networks of *Mycobacterium tuberculosis*"

**List of supplementary files:**

1. **Supplementary figures:** Figure S1 to Figure S4
2. **Supplementary Table S1:** Gene expression compendium used for developing EGRIN models.
3. **Supplementary Table S2:** EGRIN and EGRIN P-D regulatory networks.
4. **Supplementary Table S3:** Details of combinatorial regulation in regulatory network.
5. **Supplementary Table S4:** Essential and non-essential predictions from PROM, IDREAM and PRIME for glycerol and cholesterol.
6. **Supplementary Table S5:** Drug treated transcriptomes after processing through DESeq2
7. **Supplementary Table S6:** Essential and non-essential predictions from PRIME for isoniazid predictions.
8. **Supplementary File S1:** Input files for PRIME used in this study (glycerol, cholesterol and INH models; gene expression compendium; beta values) - .mat file to use with MATLAB.

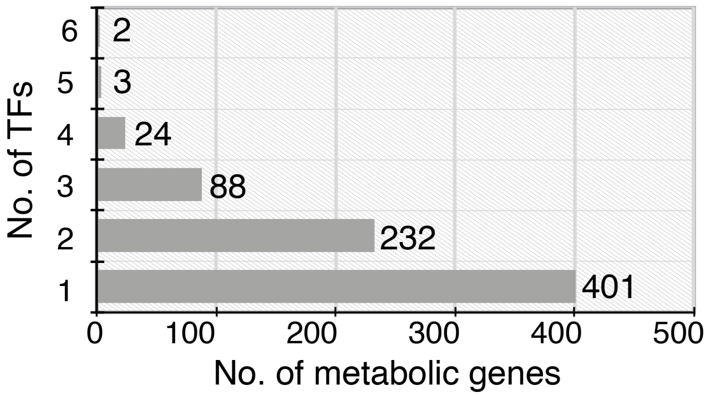


**Figure S1.** Breakdown of numbers of metabolic genes based on the number of TFs implicated in their regulation.

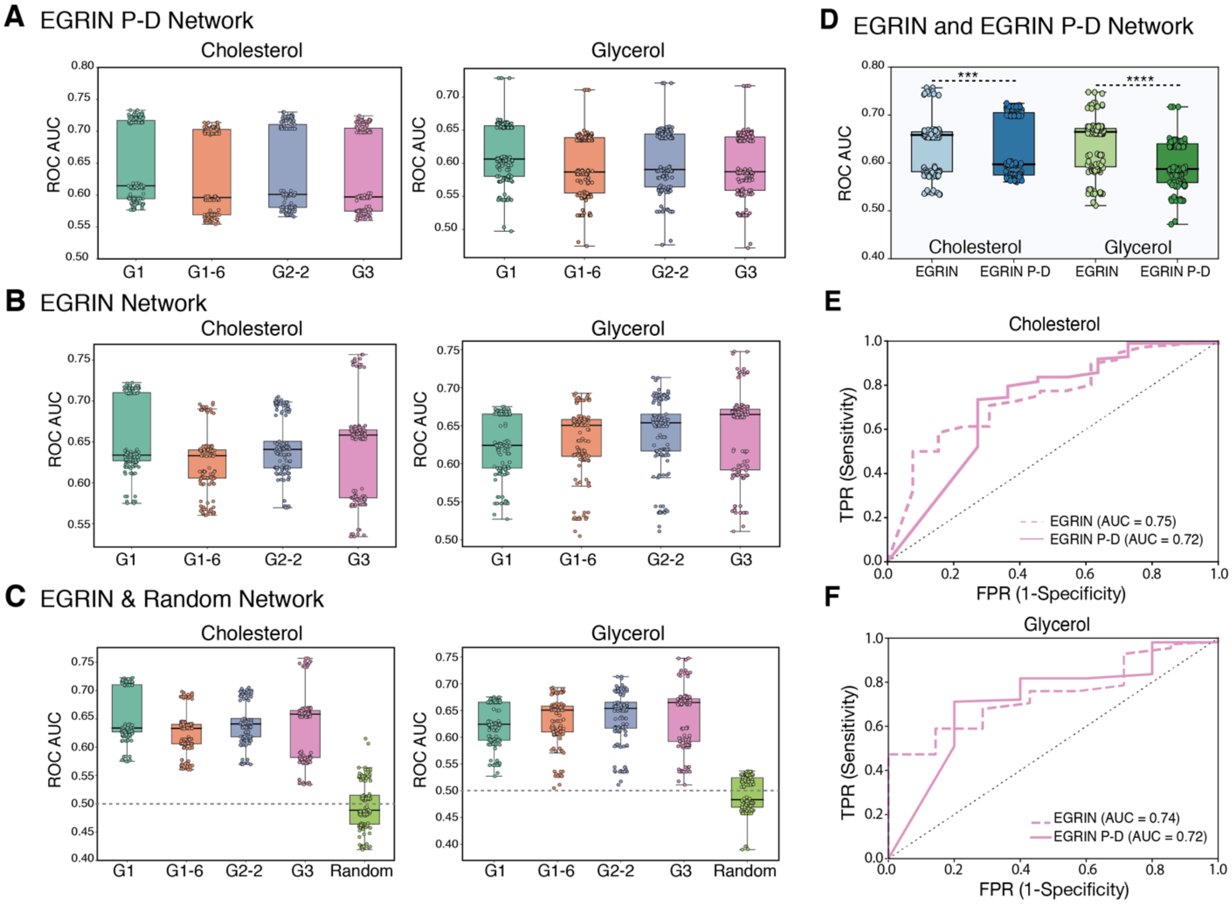


**Figure S2.** Multiple EGRIN models were constructed, based on different values assigned to the "*g*" function in the Inferelator algorithm that defines how much of the prior weights has to be assigned while retrieving the network (see Equation 2 in Bonneau et al 2006 for more details). We generated networks for four values of *g* (1.1, 1.6, 2.2, and 3). The network provided in **Table S2** has a *g* value of 3 and is the network used throughout the study. **A.** Network performance for EGRIN P-D network with all *g* values. **B.** Network performance for EGRIN network with all *g* values. **C.** Comparison of EGRIN performance with all *g* values and a random network. **D.** Comparison of performance between EGRIN and EGRIN P-D network in both (cholesterol and gycerol) growth conditions. Statistical significance was calculated as *p*-value with two sample t-test. *** p-value < 0.001 and **** p-value < 0.0001. ROC curves for EGRIN and EGRIN P-D comparison in **E**. cholesterol and **F**. glycerol growth conditions.

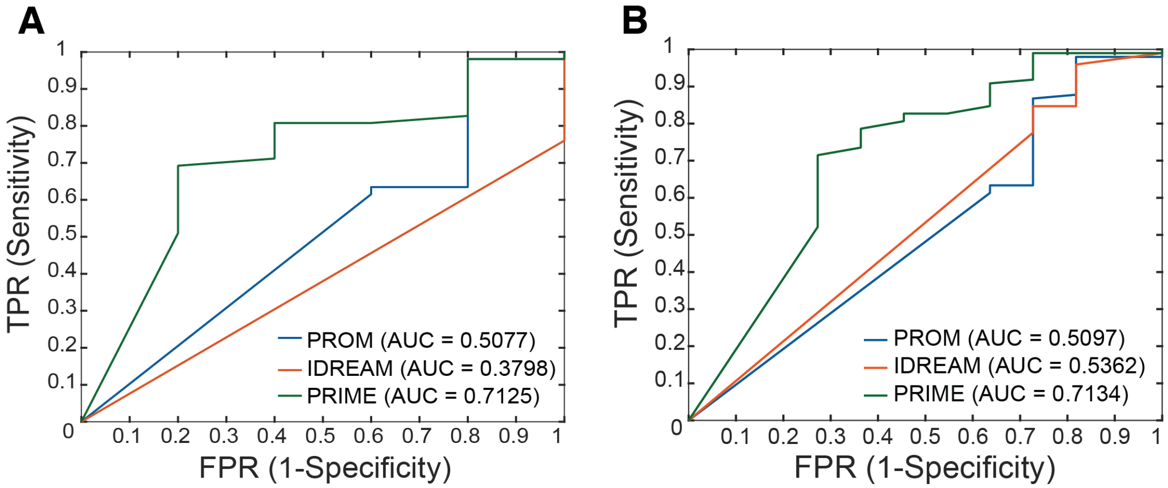


**Figure S3: ROC curve analysis for cholesterol and glycerol.** Sensitivity and specificity curve for model predictions in **A.** cholesterol and **B**. glycerol.

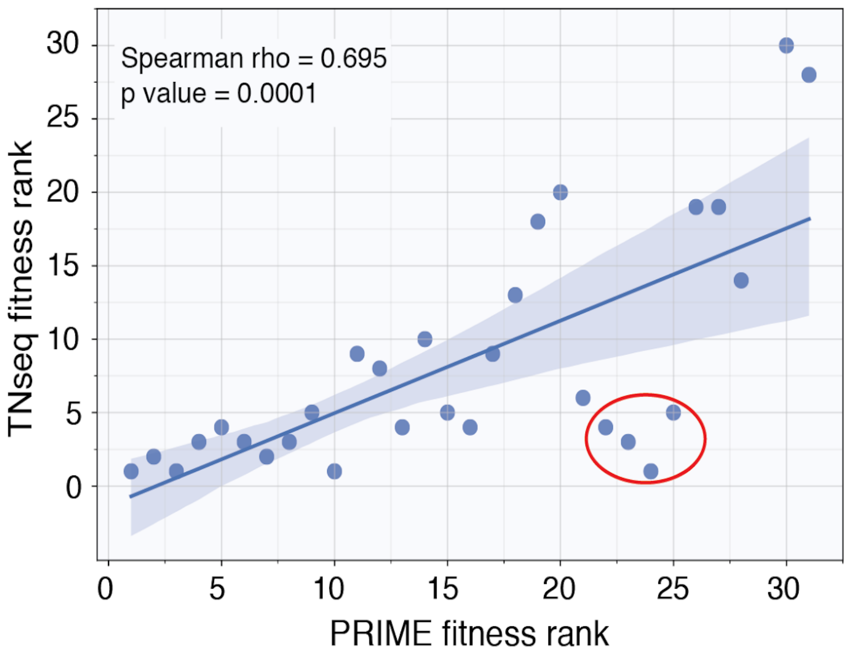


**Figure S4: TnSeq and PRIME rank correlation.** TF fitness from TnSeq experiments were compared with PRIME TF knockout fitness. The red circle highlights TFs that have <10% of target genes as part of metabolic model. These TFs were removed in **Figure 4c**.
